# Tissue-specific distribution of eggs in the definitive host drives transcriptomic and behavioral differences in *Schistosoma mansoni* miracidia

**DOI:** 10.1101/2025.08.15.670345

**Authors:** Sophie Willett, Sonja A. Olson, Rachel V. Horejsi, Chase N. Nelson, Nicolas J. Wheeler

## Abstract

Schistosomiasis is a neglected tropical disease caused by human-infective schistosomes (Trematoda: *Schistosoma*). Intestinal schistosomiasis in sub-Saharan Africa and the Neotropics is caused primarily by *Schistosoma mansoni* and is transmitted by several *Biomphalaria* planorbid snail species. Adult male and female parasites in the definitive mammalian host pair and reside in the mesenteric vasculature; females lay eggs that traverse the intestinal wall to be excreted, but a significant proportion become trapped in host tissues, especially the liver, eliciting granulomatous immune responses that underlie most disease pathology. *S. mansoni* is the primary lab model for research and, due to the abundance and ease of harvesting, liver-derived eggs are almost exclusively used to maintain the life cycle and to study miracidia and subsequent larval stages. However, recent evidence shows that eggs from the liver or intestine have key morphometric, transcriptomic, and antigenic differences, which can profoundly affect experimental outcomes. To determine whether these differences extend to the miracidia stage, we compared miracidia hatched from mouse liver and intestine-derived eggs, sequencing their transcriptomes and assessing their unstimulated behaviors over time in an arena allowing for high-resolution tracking of miracidia behavior at a large spatiotemporal scale. We found that while transcriptomic profiles of miracidia are distinguishable based on egg tissue origin, only a small subset of genes is differentially expressed. Further, basic, unstimulated behavior of miracidia that developed in different niches of the definitive host was significantly different. These different behavioral programs may reflect intrinsic developmental programming or differential viability and hardiness related to tissue origin. These findings underscore the importance of egg source in experimental design and interpretation, with significant implications for the maintenance of laboratory life cycles and the use of miracidia in schistosomiasis research.

## INTRODUCTION

The neglected tropical disease schistosomiasis affects over 250 million people around the world. Schistosomiasis is prevalent within tropical and subtropical areas of sub-Saharan Africa, South America, and eastern/southeastern Asia, especially in poverty-stricken areas with limited access to clean water or sanitation practices. *Schistosoma mansoni* is one of six species in the genus *Schistosoma* that spread this disease and is the primary lab model for all schistosomes (McManus, Dunne, Sacko, Utzinger, Vennervald, and Zhou, 2018).

Prior to becoming infective to humans as cercariae, the larval miracidia infect a snail intermediate host that inhabits freshwater environments. Cercariae infective to humans are shed from snails into aquatic surroundings where they can penetrate human skin. Cercariae will lose their tail during penetration of the definitive host and advance to the stage of schistosomulae. These schistosomulae will eventually pair in the mesenteric vasculature as male and female adults, and the females will lay eggs. Eggs will travel against the flow of blood, and mature eggs cross the intestinal wall to be excreted, thus allowing the cycle of infection to continue. However, in both humans and rodent models, many eggs will travel with the flow of blood to the liver, where they become lodged and cause granuloma formation and hepatomegaly, characteristics of progressed stages of intestinal schistosomiasis.

Eggs that travel through the intestine are excreted to hatch as miracidia that will infect the intermediate snail host. In contrast, the liver is a dead end for schistosome eggs. This presents a challenge when utilizing *S. mansoni* in the laboratory by cycling through rodents and *Biomphalaria glabrata* snails. Historically and contemporarily, liver-derived eggs are usually used to maintain the life cycle and study the behaviors of miracidia and snail-parasite interactions (DeWitt, 1955; Kagan and Geiger, 1965; Lewis, Stirewalt, Souza, and Gazzinelli, 1986; Lombardo, Pasche, Panic, Endriss, and Keiser, 2019). Intestine or feces-derived eggs are not as easy to harvest and are much less abundant resulting in less research utilizing these eggs.

Recent studies have shown that mature *S. mansoni* eggs from rodent liver or intestine have key morphometric, transcriptomic, and antigenic differences which impact experimental outcomes that use eggs as the primary point of investigation (Peterková, Konečný, Macháček, Jedličková, Winkelmann, Sombetzki, and Dvořák, 2024). However, it is unclear if these differences endure in the miracidia and later stages. Variation in the transcriptomes, behavioral differences, or other contrasts between liver and intestine miracidia may have significant effects on best-practices for life cycle maintenance or investigations of snail-schistosome interactions. We sought to understand whether the miracidia hatched from liver or intestine-derived eggs show the same contrasting characteristics as seen in the eggs.

## METHODS

### Parasite source, harvesting, and morphometrics

Female Swiss outbred mice (The Jackson Laboratory, Bar Harbor, Maine) were infected with *Schistosoma mansoni* (NMRI), and liver and intestines were harvested at the Biomedical Research Institute (BRI) (Rockville, MD) at six weeks post-infection. The infected tissues were shipped overnight in perfusion fluid to UWEC. Miracidia were harvested as previously described (Horejsi, Nelson, Ruyter, Gensch, Maasz-Seawright, Weber, Willett, Olson, and Wheeler, 2025). Briefly, tissues were homogenized in 1.2% saline and washed three times. The homogenate was transferred to a 1 L volumetric flask and filled with artificial pond water (APW; 0.77 µM FeCl_3_, 290.5 µM CaCl_2_, 207.7 µM MgSO_4_, 312.29 µM KH_2_PO_4_, 13.98 µM (NH_4_)_2_SO_4_ at pH 7.2). A light source and warm APW (32-34°C) were added to top neck of the flask. After a wait time of 15 minutes hatched miracidia that had swum to the surface were extracted for use in experiments.

Morphometrics of miracidia from three different independent harvests were measured in a 96-well plate. Wells were initially filled with 20% ethanol. Approximately 1 miracidium in APW was individually pipetted into each well. Once miracidia were added, additional ethanol was added for a penultimate concentration of 50% ethanol. After a five-minute wait time a final addition of ethanol was added for a final concentration of 70% ethanol. Miracidia were imaged inverted with an Echo Revolve (San Diego, CA USA) at 20X. Images were annotated and measured using the ECHO Pro app (v6.4.2).

### RNA extraction and sequencing

After harvesting eggs from livers and intestines and hatching in separate flasks, miracidia were maintained in APW for 1-3 hours, after which they were transferred to a 1.5 mL tube, pelleted, and APW was removed to 100 µL. 900 µL of Trizol (ThermoFisher, Waltham, MA) was added, and tubes were flash frozen in liquid nitrogen and stored at −80°C until RNA extraction. Four biological replicates (batches of paired mouse tissues) were performed on miracidia from livers and intestines.

RNA was extracted at the Gene Expression Center of the University of Wisconsin Biotechnology Center. Samples were homogenized by a TissueLyser (Qiagen, Hilden, Germany) with a 5 mm stainless steel bead in a 2 mL Safe-Lock tube (Eppendorf, Hamburg, Germany). Samples were lysed at 30 Hz for 2 min for 1 cycle and rested at on ice for 5 minutes. Chloroform was added to the samples, and an organic extraction was performed, followed by clean up using a Qiagen RNeasy Micro kit. RNA was treated on the column with DNase. Quantity and quality of purified RNA were assessed with a NanoDrop One (ThermoFisher), Qubit (ThermoFisher), and 4200 Tapestation (Agilent, Santa Clara, CA).

The NEBNext Ultra II kit was used to prepare sequencing libraries (New England Biolabs, Ipswich, MA). 10 ng of total RNA was used as input for the NEBNext Poly(A) mRNA Magnetic Isolation Module. dsDNA fragments with adapters were amplified via 17 PCR cycles and the library was purified with 0.8X NEBNext sample purification beads. Quantity and quality of libraries were assessed with a Qubit and 4200 Tapestation. All 8 libraries were barcoded, pooled, and sequenced on a full 10B lane of a NovaSeq X with a 2×150 format (Illumina, San Diego, CA).

### RNA-seq analysis

The entire RNA-seq pipeline can be found as a Snakemake workflow available on GitHub (https://github.com/wheelerlab-uwec/miracidia-distribution-ms/). RNA-seq reads were mapped to v10 of the reference genome and annotations of *S. mansoni* (Wormbase ParaSite v19) (Howe, Bolt, Shafie, Kersey, and Berriman, 2017; Buddenborg, Tracey, Berger, Lu, Doyle, Fu, Yang, Reid, Rodgers, Rinaldi, et al., 2021). FastQC (v0.12.1) and multiqc (v1.17) were used for quality control of reads before and after trimming and filtering (Ewels, Magnusson, Lundin, and Käller, 2016; “Babraham Bioinformatics - FastQC A Quality Control tool for High Throughput Sequence Data,” n.d.). Based upon the categories scored by FastQC, the following cutadapt (v4.9) parameters were used for filtering and trimming: reads were set at minimum to be 50 base pairs in length (-m 50), the first 10 bases were removed (-u 10), bases with quality scores less than 20 were removed (-q 20), NovaSeq-specific quality trimming was performed (--nextseq-trim=20), and Illumina adapter sequences were removed (Martin, 2011).

STAR (v2.7.10a) was used to perform a two-pass alignment to the reference genome (Dobin, Davis, Schlesinger, Drenkow, Zaleski, Jha, Batut, Chaisson, and Gingeras, 2013). Specific parameters for generating the genome suffix array included --sjdbOverhang=139 and --genomeSAindexNbases=13. The BAM files were sorted by coordinate and unmapped reads were included within the main SAM file. Samtools (v1.6), Picard (v3.2.0), and Qualimap (v2.3) were used to assess alignment quality control (Li, Handsaker, Wysoker, Fennell, Ruan, Homer, Marth, Abecasis, and Durbin, 2009; Okonechnikov, Conesa, and García-Alcalde, 2016; “Picard Tools - By Broad Institute,” n.d.). Duplicate reads were located with MarkDuplicates, and featureCounts (v2.0.6) was used to record counts from the BAM files (Liao, Smyth, and Shi, 2014). Specific parameters used were: -p, --countReadPairs, --fraction, --ignoreDup, and-M to account for paired-end reads and multi-mapped reads.

Read counts were transformed and normalized, and differentially expressed genes (DEGs) were identified using the R package DESeq2 (v1.48.1) (Love, Huber, and Anders, 2014). PCA was performed on transformed counts and compared based on the tissue type the reads originated from. Differentially expressed genes were identified using a p-adjusted value of <0.05 and log2FoldChange values of >1 or <-1.

### Miracidia behavioral experiments

Recording miracidia ethology was done using custom designed and assembled basic ethology arenas (Supplemental Figure 1), as previously described (Horejsi, Nelson, Ruyter, Gensch, Maasz-Seawright, Weber, Willett, Olson, and Wheeler, 2025). Each section of the arenas was created from 1.6 mm (1/16”) thick clear acrylic (McMaster-Carr, Elmhurst, IL USA) and laser-cut with a Trotec Speedy 360 80-Watt laser CO2 engraver/cutter (Marchtrenk, Austria). Once acrylic pieces were cut, each was cleaned with 70% ethanol and dried completely to remove any remaining dust or contaminants from the laser cutting process. Three clear acrylic pieces were cut out for the basic ethology arenas: a lid with a thin etched groove border and a central pore to load the parasites, a frame, and a base piece with the same thin etched groove border to align the frame and lid. To secure the acrylic pieces together, methyl ethyl ketone (MEK) was applied using a micropipette. Finished fabricated basic ethology arenas were closed with tape over pore opening to stop contamination of dust from entering arenas during storage.

**Figure 1.**
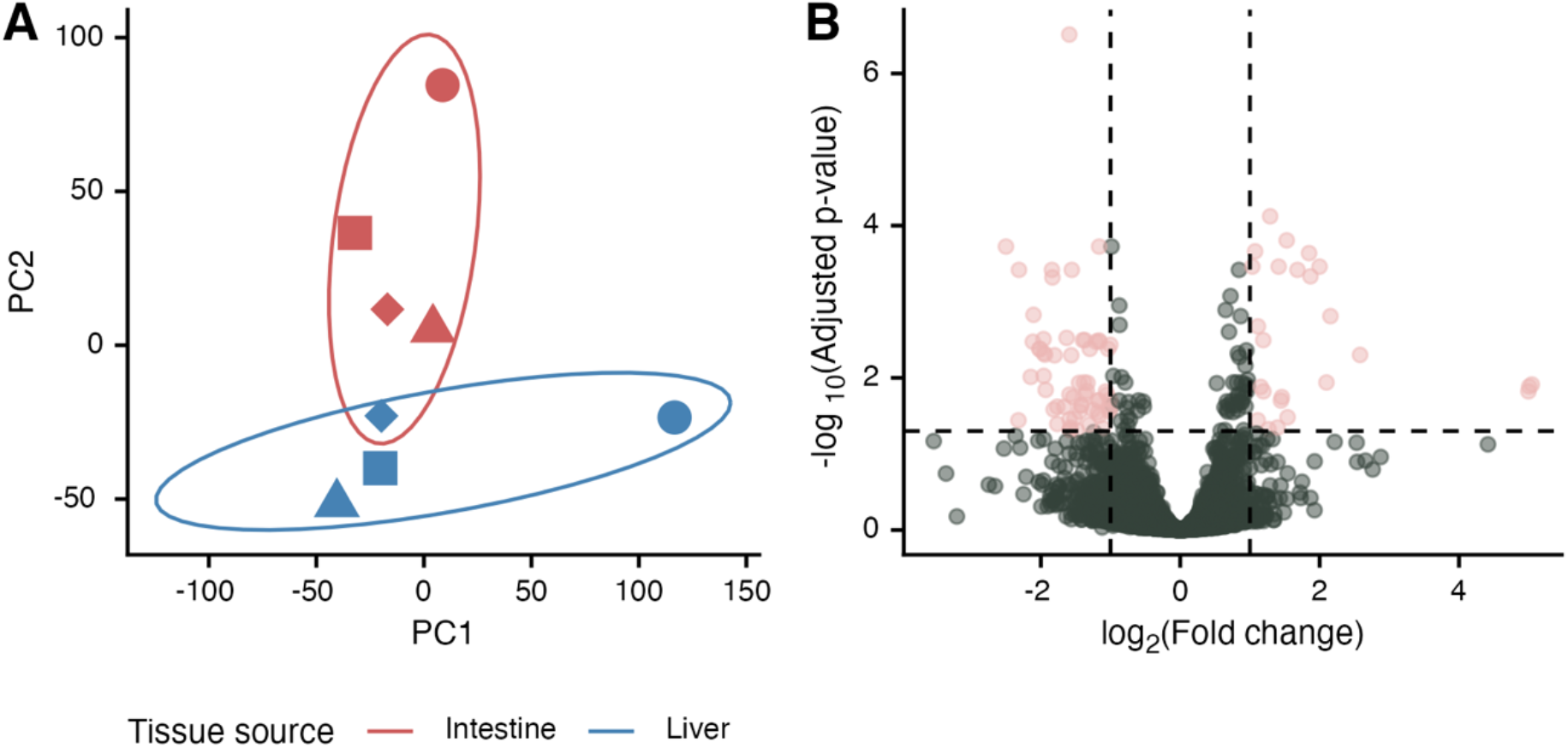
RNA sequencing of *Schistosoma mansoni* miracidia hatched from eggs extracted from mouse livers or intestines. (A) PCA plot of transformed counts that depicts clusters based on tissue source. Point shapes correspond to biological replicates (miracidia hatched from eggs harvested from livers/eggs from the same mice). Ellipses represent a 68% confidence region (1 standard deviation) for the bivariate normal distribution of the plotted points. (B) Volcano plot showing differentially expressed genes (DEGs, pink). DEGs have a statistical significance of p-adjusted ≤ 0.05 (horizontal dotted line) and a |log_2_ fold-change| > 1 (vertical dotted lines).

For high-definition video imaging, we used the InVision (<u>in</u>vertebrate <u>vision</u>) for tracking parasite movements and behaviors (Horejsi, Nelson, Ruyter, Gensch, Maasz-Seawright, Weber, Willett, Olson, and Wheeler, 2025). Before imaging, basic ethology arenas were prepped by cleaning with 70% ethanol and filled with APW using a micropipette. Calculations of miracidia density was conducted from each tissue source (liver or intestines) by counting five 10 µL aliquots of miracidia stained with 10 µL of 1:5 diluted Lugol’s iodine. Dilutions were performed to create a final density of 1 miracidium/µL to use in ethology assays. 20 µL of APW was removed from pre-filled arenas and replaced with 20 µL of miracidia (>15 minutes old). Some harvests had reduced yields requiring manual retrieval of miracidia; in these cases, live miracidia were manually pipetted from a Petri dish under a Zeiss Stemi 305 microscope (Oberkochen, Germany) into basic ethology arenas. Approximately 20 miracidia were pipetted per arena. Arenas were then transferred to InVision stage for video recordings. Five biological replicates were performed with miracidia used from separate infections. Two replicates were conducted with 1 hour video recordings, and the other three replicates were conducted with 4 hour long video recordings. All five recordings were taken at 15 frames per second (FPS). Four basic ethology arenas were used per experiment – two arenas per tissue source.

Miracidia were tracked in videos using the invision-tools (https://github.com/wheelerlab-uwec/invision-tools), as previously described (Horejsi, Nelson, Ruyter, Gensch, Maasz-Seawright, Weber, Willett, Olson, and Wheeler, 2025). Tracks were split into 5-second chunks, and 43 quantitative features were extracted from each chunk. Each feature was modeled as a linear mixed effect model (feature ~ tissue * frame_start + (1 + date/track)) using nlme where the fixed response variables were the tissue source and the beginning frame number for the track chunk. The interaction between the tissue source and the frame start was also analyzed, and correlation between chunks was controlled for with corAR1 of the nlme package (Pinheiro and Bates, 2025). P-values were adjusted with the Benjamini & Hochberg correction, and the false discovery rate was set to 0.05 (Benjamini and Hochberg, 1995). Effect sizes were calculated using Cohen’s D using the pooled standard deviation for each feature (Cohen, 1988).

### Data availability

All analytical code is available on a public GitHub repository (https://github.com/wheelerlab-uwec/miracidia-distribution-ms). RNA-seq read data is available for download via the NCBI SRA (BioProject ID PRJNA1299362).

## RESULTS

### Miracidia derived from eggs from different tissue sources are transcriptomically distinct

It was previously reported that schistosome eggs harvested from different host tissues were transcriptomically distinct (Peterková, Konečný, Macháček, Jedličková, Winkelmann, Sombetzki, and Dvořák, 2024). To test whether these differences endured in miracidia, we hatched miracidia from liver and intestines and allowed them to equilibrate for 1-3 hours at room temperature before extracting RNA to be sequenced.

Global transcriptomic differences between the respective samples were distinguished based upon their originating tissue type (Figure 1A). The clustering of each was heavily influenced by the tissue type the sample originated from. This result suggests that the miracidia hatched from the different tissue types are transcriptomically distinct.

Clustering accounts for global transcriptomic differences, and we next analyzed genes that were differentially expressed between miracidia derived from different host tissues. We hypothesized that some differentially expressed genes (DEGs) between liver miracidia/intestine miracidia would be shared with previously identified DEGs between mature liver eggs/intestine eggs. We also hypothesized that there would be fewer DEGs overall. Mature eggs derived from either the liver or intestine were shown to have 1339 DEGs (Peterková, Konečný, Macháček, Jedličková, Winkelmann, Sombetzki, and Dvořák, 2024), while we only detected 87 DEGs in equilibrated miracidia (Table 1, Figure 1B). Of these 87, 28 of them (32%) were shared with the DEGs from liver/intestine mature eggs. Differences in number of DEGs are likely in part due to technical differences in library preparation and sequencing, but stark reduction in liver/intestine derived miracidia may also be due to the synchronization of the transcriptomes as the parasites get temporal distance from their distinct developmental niches as eggs. These data show that miracidia derived from eggs hatched from host livers or intestines are transcriptomically distinct but that they have relatively few DEGs when compared to eggs harvested directly from host tissues.

### Miracidia derived from eggs from different tissue sources have negligible differences in morphometric features

To begin to investigate whether transcriptomic differences had emergent morphological or behavioral effects, we measured the area and length of miracidia hatched from eggs derived from liver and intestine tissues (Figure 2). There was no significant difference in area (5751.12 µm^2^ liver miracidia vs. 5794.47 µm^2^ intestine miracidia), but there was a significant difference in length (143.75 µm liver miracidia vs. 136.99 µm intestine miracidia). However, the size of that difference was trivial (4.7%). These findings suggest miracidia hatched from eggs originating from different tissues have miniscule morphometric differences.

**Figure 2.**
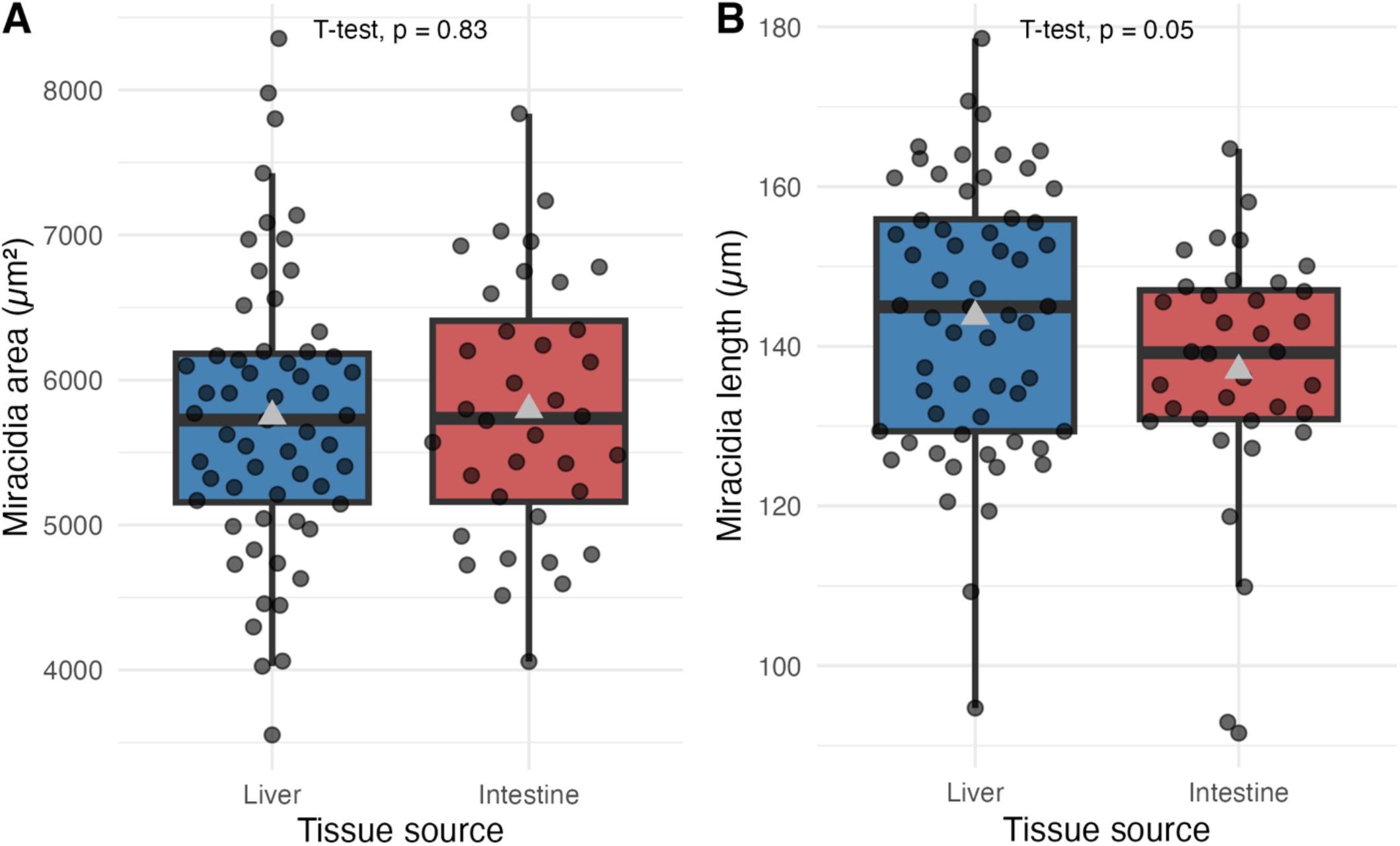
Measurements of *Schistosoma mansoni* miracidia hatched from eggs extracted from mouse livers or intestines. Boxplots representing area (A) and length (B) of miracidia obtained from liver and intestinal tissues. The mean is represented by a gray triangle.

### Miracidia derived from eggs from different tissue sources have distinct behavioral programs

We next examined the behavior of miracidia hatched from eggs derived from liver and intestine tissues. Parasites were recorded and tracked in basic ethology arenas for one hour, and 43 behavioral features were extracted from the resulting track data. Linear mixed models revealed that the fixed effect of originating tissue produced significant differences in 33 out of 43 features, with 15 of these also showing significant interaction effects (Figure 3A). The direction of the effect differed across feature categories. For example, speed tended to be higher in miracidia from liver eggs while heading variance (how much a parasite’s direction changes over time) and angular velocity metrics were higher in miracidia from intestine eggs (Figure 3B/C). Overall, miracidia from liver eggs tended to be faster and swim straighter, while miracidia from intestine eggs were slower and had curvier paths, and the aggregated behavioral profiles suggest distinct behavioral programs. Principal component analysis (PCA) of the feature summaries demonstrated distinct clustering of liver and intestine-derived miracidia, with minor overlap due to a single outlier sample from the intestine group (Figure 3D).

**Figure 3.**
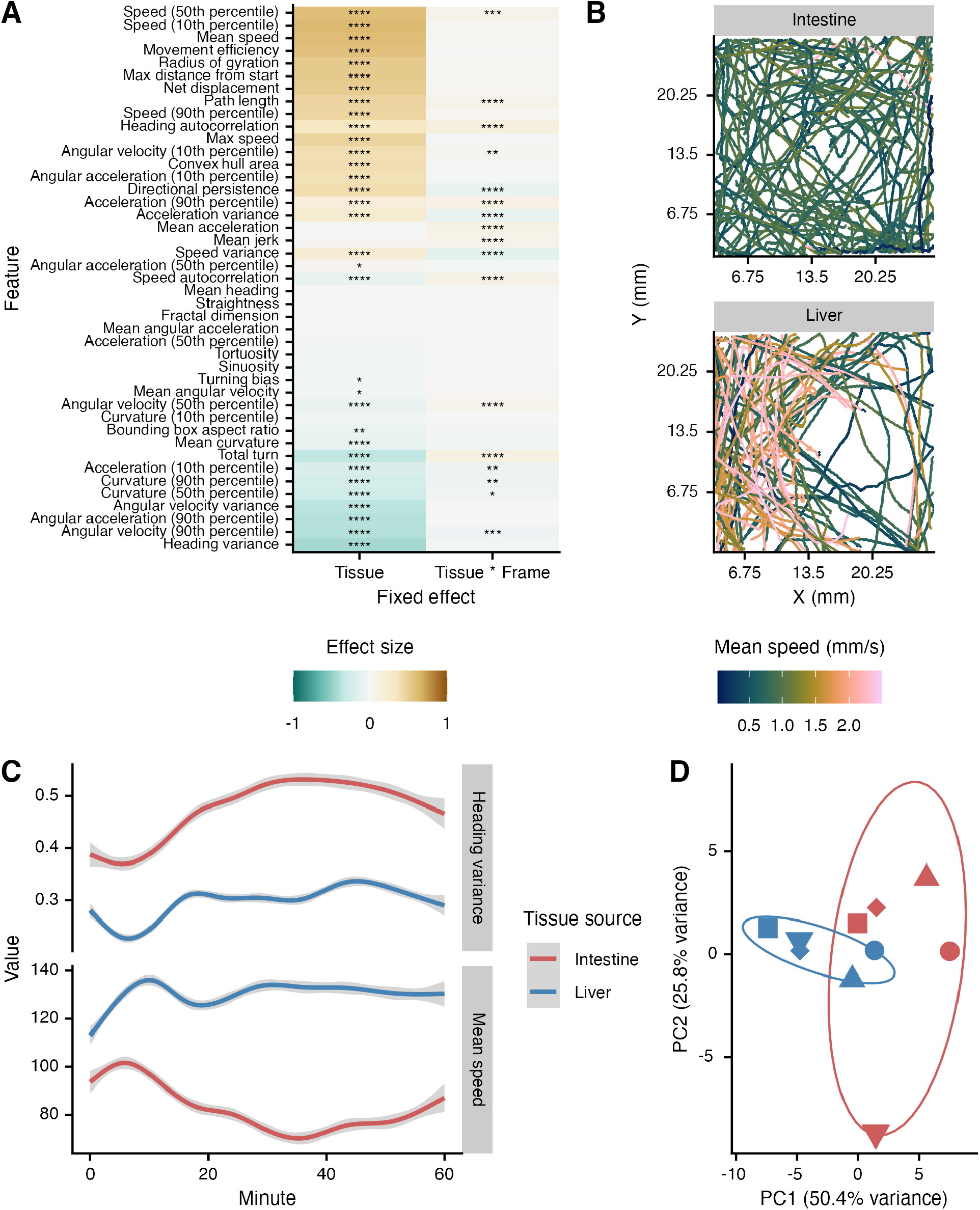
Base behavior differs between *Schistosoma mansoni* miracidia derived from different tissue locations. (A) 43 behavioral features were extracted from chunks of tracks recorded in basic ethology arenas. Main effect sizes for tissues were compared in addition to the interaction between the frame number and tissue. Colors represent standardized effect size (Cohen’s D) with liver miracidia as the reference. Brown indicates a larger positive (the value is greater in liver miracidia) and blue signifies a larger negative effect (the value is smaller in liver miracidia). Significant differences between behavioral features are represented by asterisks (*p < 0.05; **p < 0.01; ***p < 0.001; ****p < 0.0001). (B) Representative sample of tracks from one biological replicate (miracidia hatched from eggs harvested from livers/eggs from the same mice). Mean speed was calculated for 5-second chunks. (C) Representative features illustrating effects caused by tissue origination and its interaction with the timepoint. (D) Principal components analysis using means from the features shown in (A). Point shapes correspond to biological replicates. Ellipses represent a 68% confidence region (1 standard deviation) for the bivariate normal distribution of the plotted points.

Miracidia from both tissue sources showed substantial attrition and variation in hardiness over 4-hour long videos (Figure 4). In general, videos of miracidia originating in the liver had more initial observations, corresponding to more miracidia in the arena, than videos of miracidia from the intestine. The number of observations decreased sharply for both categories of miracidia due to death and/or depletion of energy. Interestingly, one sample of miracidia hatched from intestine-derived eggs had a much smaller negative slope, demonstrating the substantial batch variation. Only 3 of 8 videos still had moving miracidia after 3 hours of recording.

**Figure 4.**
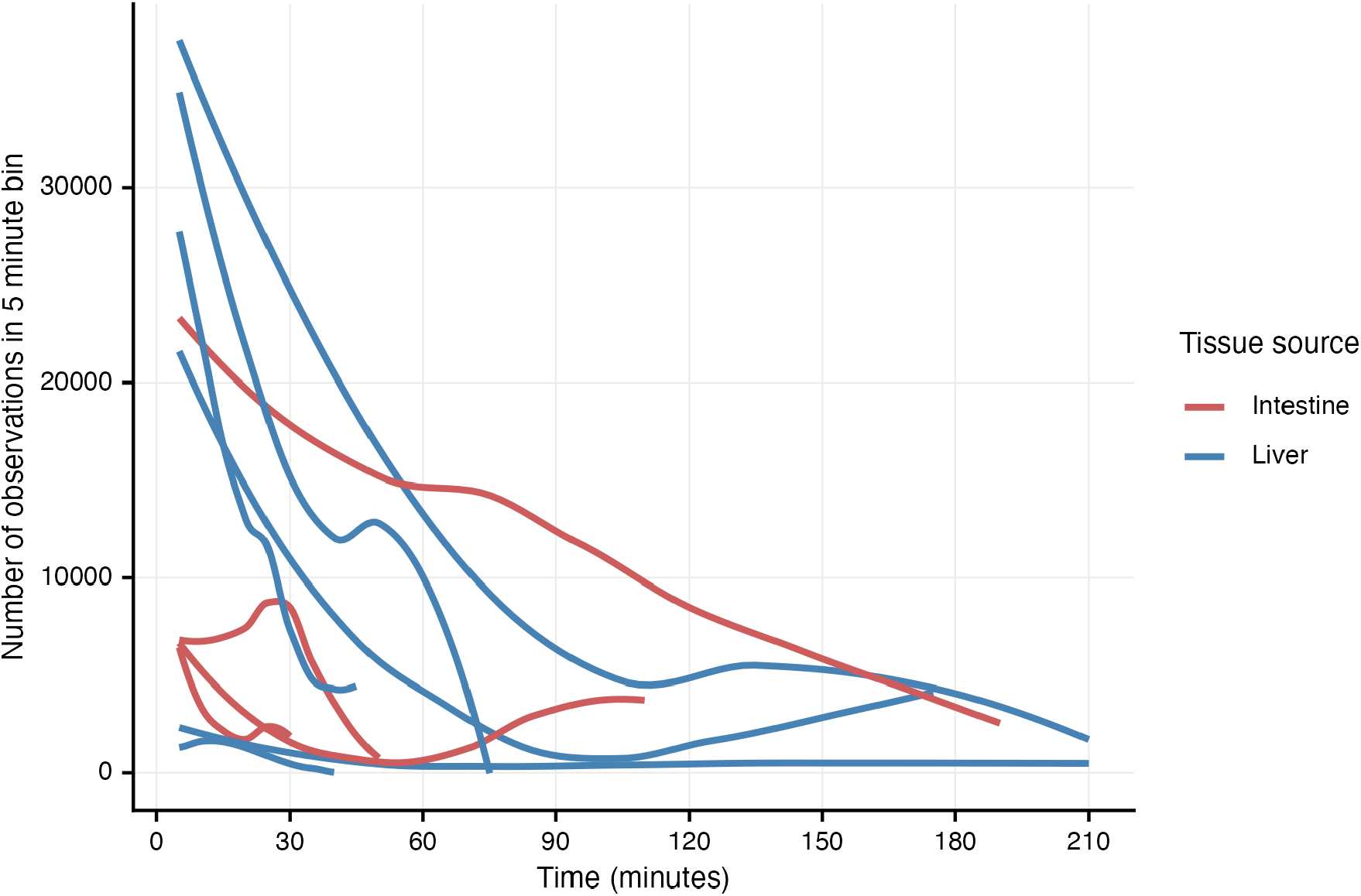
Attrition of *Schistosoma mansoni* miracidia over 4-hour long videos. The number of observations (i.e., detected miracidia) over 5-minute video chunks were tallied. Lines connecting data points were smoothed using LOESS fitting. Lines represent technical replicates (separate arenas) from 4 biological replicates. Some arenas had no tracks and thus are not shown.

## DISCUSSION

The miracidia of *Schistosoma mansoni* have long been used as a model species in the field of behavioral parasitology and the study of snail-trematode interactions, and it is a critical choke point in the life cycle of the species that cause schistosomiasis (Saladin, 1979). Much contemporary research is conducted using miracidia obtained from the host’s liver tissue (Spaan, Pennance, Laidemitt, Sims, Roth, Lam, Rawago, Ogara, Loker, Odiere, et al., 2023; Archer, Yeo, Gadd, Pennance, Cunningham, Juhàsz, Jones, Chammudzi, Kapira, Lally, et al., 2024; Attenborough, Rawlinson, Diaz Soria, Ambridge, Sankaranarayanan, Graham, Cotton, Doyle, Rinaldi, and Berriman, 2024; Poteaux, Ripoll, Sarrazin, Blanchard, Guillou-Duvoid, Gourbal, Hirbec, and Duval, 2024), and snail/schistosome core facilities often use liver-derived miracidia for snail infections to maintain the life cycle (Lewis, Stirewalt, Souza, and Gazzinelli, 1986; “Exposure of snails to miracidia - Biomedical Research Institute,” 2022). This is due to the high abundance of mature eggs in rodent liver and its ease of harvest, a fact that was experienced in this study as well. The reality of working with miracidia harvested from intestinal tissues is that harvests frequently yielded low counts, making it a challenge for loading enough healthy, viable miracidia into ethology arenas. At times, this required intestinal miracidia to be subjected to several centrifuge steps and frequent manual pipetting, which may lead to parasite attrition.

For these reasons, even though we have shown significant transcriptomic and behavioral differences between miracidia arising from eggs from different host tissues, continued use of miracidia obtained from liver tissues may be justified unless experimental fidelity with the natural life cycle and infection process is required. For example, some recent studies use miracidia hatched from eggs derived from human or rodent feces (Pennance, Tennessen, Spaan, McQuistan, Ogara, Rawago, Andiego, Mulonga, Odhiambo, Mutuku, et al., 2025), and this was the historical approach by other select investigators (Schreiber and Schubert, 1949; Standen, 1949, 1951). When miracidia behavior or the snail-miracidia interactions are the subject of investigation, it may be wise to consider using a parasite source that mimics the natural life cycle, especially if high numbers (i.e., thousands) of parasites are not required.

Investigation of the differences between later stages of parasites originally derived from liver and intestines should be continued to better understand how long the transcriptomic and behavioral differences endure. The fewer number of miracidia DEGs in comparison to previous studies of eggs suggest that the transcriptomes may be synchronizing with developmental progression (Figure 1, Table 1) (Peterková, Konečný, Macháček, Jedličková, Winkelmann, Sombetzki, and Dvořák, 2024). Whether transcriptome synchronization is complete after snail infection, cercarial shedding, or even later remains to be seen. Regardless, even with fewer transcriptomic differences, emergent behavioral phenotypes remain (Figure 3). It will be important to understand continued divergence or lack thereof in behavior or antigen availability during parasitization of the intermediate and definitive hosts. Indeed, it may be that epigenetic effects initiated during egg development persevere even to adulthood or the next generation, and these could have a variety of meaningful effects on experiments that use mature parasite stages or best practices in parasite husbandry.

## ACKNOWLEDGMENTS

*Schistosoma mansoni*-infected mouse tissue was provided by the Schistosomiasis Resource Center of the Biomedical Research Institute (Rockville, MD) through NIH NIAID Contract HHSN272201700014I. Funding was provided by Student Blugold Commitment Differential Tuition funds through the University of Wisconsin-Eau Claire and the UWEC Office of Academic Affairs. Some computational resources used for this study were provided by the Blugold Center for High-Performance Computing under NSF grant CNS-1920220. NJW, SEW, SAO, RVH, and CNN were supported by NIH NIAID R15 AI183095. The authors would like to thank UWEC BIOL 343 students for collaborating on the RNA-seq analysis and the UWEC Makerspace for laser-cutting services.

