## Supplementary figures and images for "Tissue-specific distribution of eggs in the definitive host drives transcriptomic and behavioral differences in *Schistosoma mansoni* miracidia"

### Supplemental Figure 1

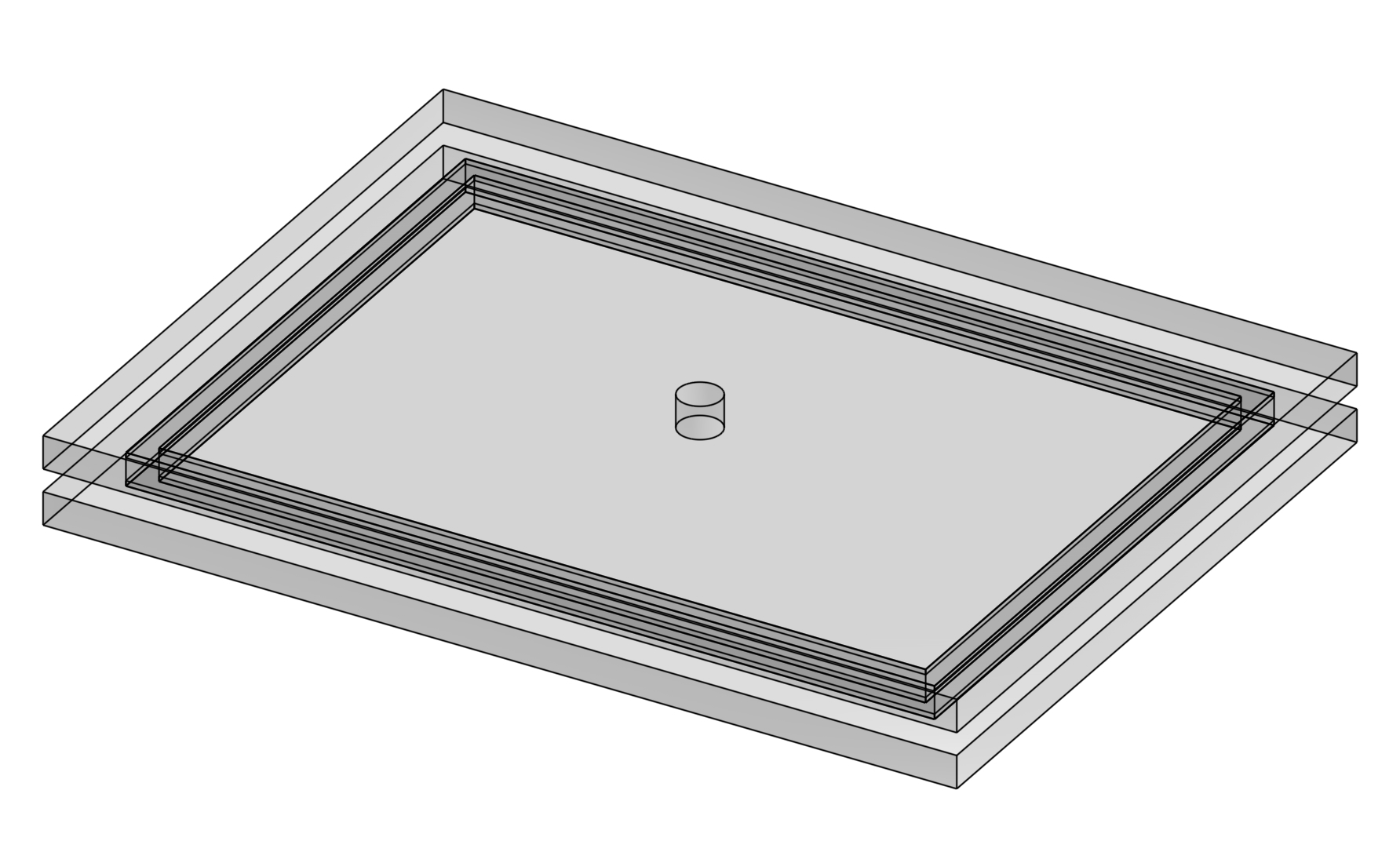
